# A widespread coral-infecting apicomplexan contains a plastid encoding chlorophyll biosynthesis

**DOI:** 10.1101/391565

**Authors:** Waldan K. Kwong, Javier del Campo, Varsha Mathur, Mark J. A. Vermeij, Patrick J. Keeling

## Abstract

The Apicomplexa are an important group of obligate intracellular parasites that include the causative agents of human diseases like malaria and toxoplasmosis. They evolved from free-living, phototrophic ancestors, and how this transition to parasitism occurred remains an outstanding question. One potential clue lies in coral reefs, where environmental DNA surveys have uncovered several lineages of uncharacterized, basally-branching apicomplexans. Reef-building corals form a well-studied symbiotic relationship with the photosynthetic dinoflagellate *Symbiodinium*, but identification of other key microbial symbionts of corals has proven elusive. Here, we used community surveys, genomics, and microscopy to identify an apicomplexan lineage, which we name ‘corallicola’, that was found in high prevalence (>80%) across all major groups of corals. In-situ fluorescence and electron microscopy confirmed that corallicola lives intracellularly within the tissues of the coral gastric cavity, and possesses clear apicomplexan ultrastructural features. We sequenced the plastid genome, which lacked all genes for photosystem proteins, indicating that corallicola harbours a non-photosynthetic plastid (an apicoplast). However, the corallicola plastid differed from all other known apicoplasts because it retains all four genes involved in chlorophyll biosynthesis. Hence, corallicola shares characteristics with both its parasitic and free-living relatives, implicating it as an evolutionary intermediate, and suggesting that a unique ancestral biochemistry likely operated during the transition from phototrophy to parasitism.

---

Apicomplexan parasites rely on highly specialized systems to infect animal cells, live within those cells, and evade host defenses. Recently, it has come to light that these parasites evolved from phototrophic ancestors; two photosynthetic ‘chromerids’, *Chromera velia* and *Vitrella brassicaformis*, were isolated from coral reef environments and were found to be the closest free-living relatives to the parasitic Apicomplexa (Moore et al. 2008; Oborník et al. 2012; Woo et al. 2015). That the photosynthetic relatives of apicomplexans are somehow linked to coral reefs has prompted a major reevaluation of the ecological conditions and symbiotic associations that drove the evolution of parasitism (Cumbo et al. 2013; Janouškovec et al. 2013; Mohamed et al. 2018; Mathur et al. 2018).

Corals (class Anthozoa) have not been traditionally considered a common host to apicomplexans: sporadic reports over the last 30 years include the morphological description of a single coccidian, *Gemmocystis cylindrus*, from histological sections of eight Caribbean coral species (Upton & Peters 1986), and the detection of apicomplexan 18S rRNA gene sequences (called “Type-N”) in Caribbean and Australian corals (Toller et al. 2002; Šlapeta & Linares 2013). Intriguingly, plastid 16S rRNA gene surveys have revealed a number of uncharacterized Apicomplexan-Related Lineages (ARLs), notably “ARL-V”, being tightly associated with reefs worldwide (Janouškovec et al. 2012; Mathur et al. 2018). They appear to occupy an intermediate phylogenetic position between that of the obligately parasitic Apicomplexa and the free-living chromerids, making them promising candidates to study how the evolutionary transition between these different lifestyles occurred.

## Results and Discussion

To address this question and to reconcile the currently incomparable data on the extent of apicomplexan diversity in corals, we sampled diverse anthozoan species from Curaçao (southern Caribbean), and surveyed the composition of both their eukaryotic and prokaryotic microbial communities (Table S1). To avoid inclusion of host DNA in the eukaryotic dataset, genes were amplified with a primer set designed to exclude metazoans (see Methods and Materials). From 43 total samples representing 38 coral species, we recovered apicomplexan Type-N 18S rRNA and ARL-V 16S rRNA genes from 62% and 83% of samples, respectively (Fig. 1a, Table S2). Type-N reads were only detected in corals that were also positive for ARL-V, suggesting they come from the same organism (the lower recovery of Type-N was likely due to the dominance of *Symbiodinium* reads in the 18S rRNA gene survey). Excluding *Symbiodinium*, Type-N was the most common microbial eukaryote detected in corals, at 2.1% of total sequence reads (56% of all non-*Symbiodinium* reads). No other apicomplexan-related lineage was present, except for 6 reads of *Vitrella* 16S rRNA in a single sample.

**Figure 1.**
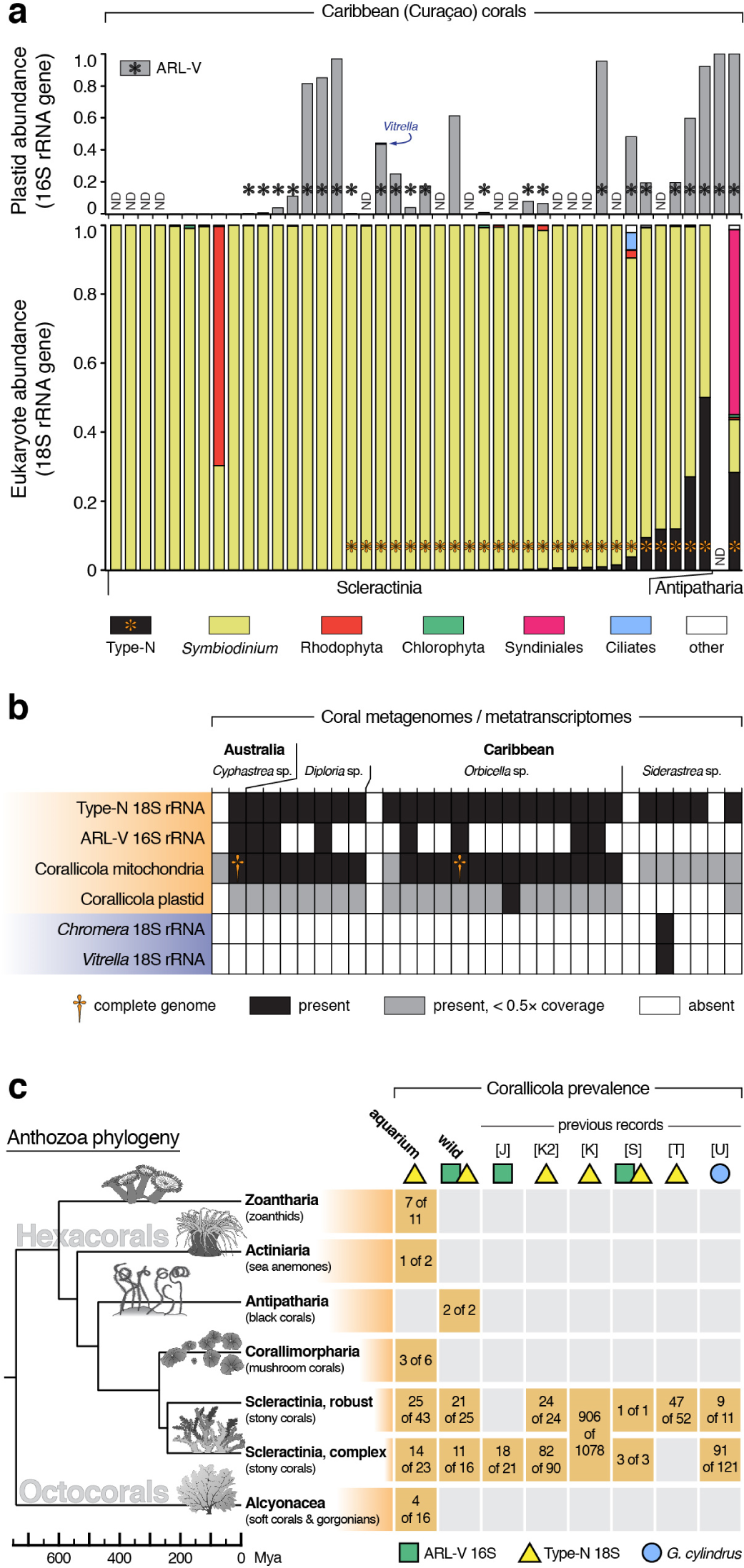
A single apicomplexan symbiont is present in diverse corals. **(a)** Relative abundance of Apicomplexa-derived plastid 16S rRNA gene sequences (top) and total microbial eukar-yote communities based on 18S rRNA gene sequencing (bottom). Each column represents data from a single coral sample. ARL-V and Type-N were the predominant Apicomplexa-related sequences (presence denoted by asterisks), and showed a correlated presence within samples. The 16S rRNA gene primers used excluded detection of *Symbiodinium* plastids. ND, not determined due to insufficient read depth. **(b)** Presence of Apicomplexa-derived sequences in public metagenomic and metatranscriptomic datasets. Host species indicated at top. Shading indicates presence or absence, or coverage of assembled contigs compared to the complete organellar genomes. **(c)** Presence of corallicola across the anthozoan phylogeny. Data includes aquarium and wild-collected samples from this study (bolded columns), and wild samples from previous studies (Upton & Peters 1986, [U]; Toller et al. 2002, [T]; Šlapeta & Linares 2013, [S]; Kirk et al. 2013, [K]; Kirk et al. 2013b, [K2]; Janouškovec et al. 2013, [J]) that used various methods to detect ARL-V, Type-N, and/or *G. cylindrus* (Table S4). Coral phylogeny was inferred from published data (see Methods and Materials).

We also searched 31 publically available coral metagenomic and metatranscriptomic datasets amounting to 15.8 Gbp of assembled sequence (Table S3). Sequences corresponding to Type-N 18S rRNA and ARL-V 16S rRNA were present in 27 and 8 datasets, respectively (Fig. 1b); the discrepancy probably reflects the lower relative copy number of plastid rRNA genes. We further identified a suite of organelle-derived protein-coding genes (see below for details), and in each case found a single apicomplexan sequence to be predominant in all samples (Fig. 1b, S1). These data are all consistent with the presence of a single dominant apicomplexan lineage in corals, for which we propose the name “corallicola”, meaning “coral-dwellers”.

The high prevalence of corallicola in wild corals is suggestive of a tight symbiosis, across a broad diversity of coral species. To test the host range of this symbiosis, we obtained 102 commercially aquacultured corals and their close relatives, representing at least 61 species from across the major clades of Anthozoa. We detected corallicola 18S rRNA genes in 53% of samples, including in soft-bodied octocorals, zoanthids, anemones, and corallimorphs (Fig. 1c). Combined with data from previous studies (Table S4), corallicola was collectively found in 1269 of 1546 samples (82% prevalence), from all parts of the anthozoan phylogeny thus far examined.

To further characterize corallicola biology, we chose an aquacultured green mushroom coral as a model (*Rhodactis* sp.; Fig. 2a). Corallicola cells were first visualized by hybridizing coral tissues with fluorescent probes specific to Type-N 18S rRNA and ARL-V 16S rRNA. Overlapping fluorescence signal from both probes was observed within the cnidoglandular lobes of the mesenterial filaments (Fig. 2b,c). This tissue region is dense with nematocysts and secretory cells, as mesenterial filaments help digest food within the gastric cavity, and may also be expelled from the polyp body for defense (Lang 1973). The only formally described apicomplexan from coral, *G. cylindrus,* was also found in mesenterial filaments (Upton & Peters 1986). No genetic sequence data for *G. cylindrus* is available, but similarities in localization, cell size (~10 µm), and Coccidia-like morphology indicate it is likely a corallicolid.

**Figure 2.**
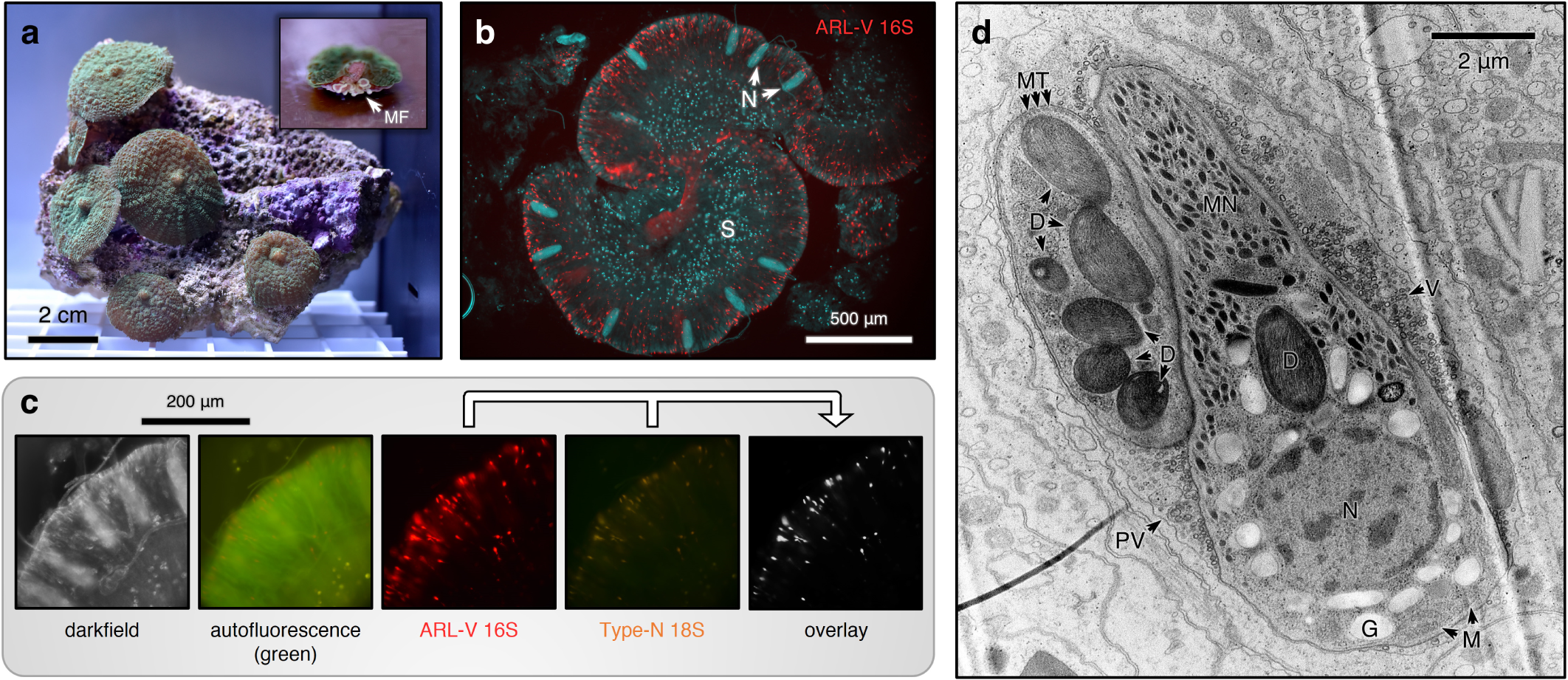
Corallicola is located intracellularly within the mesenterial filaments, and possesses cellular features distinctive of apicomeplexans. **(a)** The *Rhodactis* sp. coral from which corallicola was imaged and sequenced. Inset shows cross section with mesenterial filaments (MF) lining the gastric cavity. **(b)** Fluorescent in-situ hybridization imaging localizes corallicola (red, plastid rRNA) to cnidoglandular bands. N, nematocysts; S, *Symbiodinium* cells. **(c)** Co-localization of ARL-V (red) and Type-N (orange) signals. Unlike *Symbiodinium*, corallicola does not exhibit autofluoresence. **(d)** Transmission electron micrograph showing corallicola ultrastructure. Two adjacent cells are depicted, oriented perpendicularly. D, darkly-staining organelles; G, polysaccharide granules; M, mitochondria; MT, microtubules; MN, micronemes; N, nucleus; PV, parasitophorous vacuole; V, extracellular vesicles.

Transmission electron microscopy of infected tissue showed cells with classical apicomplexan features (e.g., a conical cortex of microtubles and inner membrane complex) residing within a parasitophorous vacuole, inside host cells (Fig. 2d). Corallicola cells were often found closely clustered together, consistent with reproductive stages in other apicomplexans (e.g., schizongony or oocyst development). Intriguingly, cells contained numerous large (up to 1.6 µm), dark-staining, elliptical organelles with striated internal features (Fig. S2). These distinctive structures could be homologous to known apicomplexan compartments such as rhoptries or even plastids, but identification will require localization of functionally relevant marker proteins.

The most fundamental question about the relationship between corallicola and their host, and by extension how they affect our views of apicomplexan origins, is whether they are photosynthetic or parasitic. To address this, we sequenced the corallicola plastid genome, and re-assessed their position in the phylogenetic tree of apicomplexans using plastid, mitochondrial, and nuclear data (Fig. 3a). We first retrieved all possible corallicola plastid and mitochondrial sequences from the metagenomic datasets using homologues from *C. velia* and *V. brassicaformis*, and from the parasitic apicomplexans *Toxoplasma gondii* and *Plasmodium falciparum*, as search queries (see above). Close matches were retrieved from 29 of 31 datasets and manually inspected to confirm their apicomplexan origin. Two complete mito-chondrial genomes were assembled, as well as fragments of plastid genomes, but insufficient plastid read coverage and the presence of sequence variants prevented an unambiguous metagenomic assembly (Fig. 1b). Thus, we used a combination of new metagenomic sequencing and primer walking to obtain a complete corallicola plastid genome from the *Rhodactis* sp. host.

**Figure 3.**
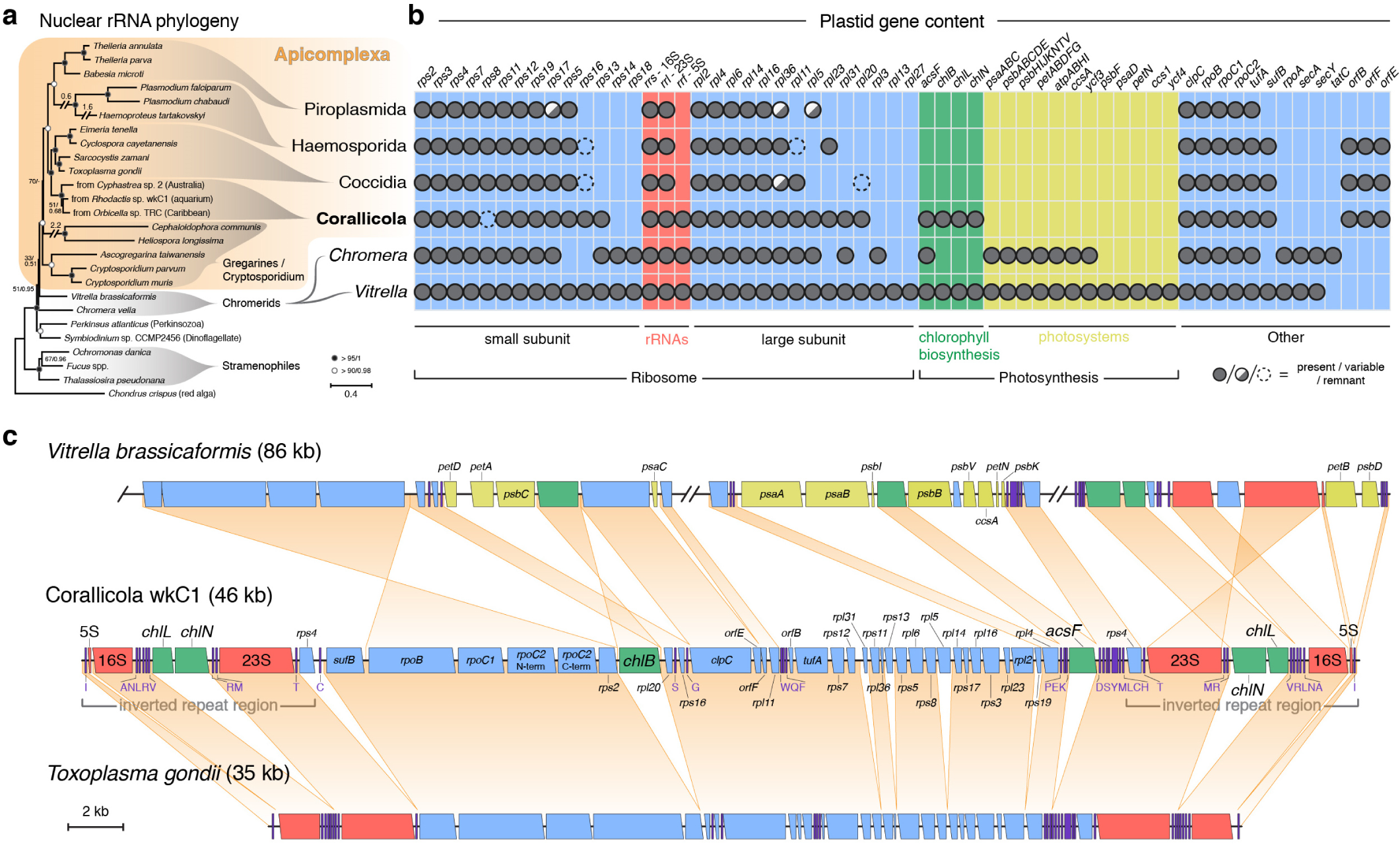
Corallicola shares traits of both the parasitic Apicomplexa and free-living photosynthetic relatives. **(a)** Maximum likelihood (ML) phylogeny based on concatenation of nucleus-encoded 18S, 5.8S, and 28S rRNA genes. Corallicola branches within the parasitic Apicomplexa, as sister to the Coccidia. Values before and after slash at nodes denote ML bootstrap values and Bayesian posterior probabilities, respectively. Accessions numbers for sequences used are listed in Table S5. **(b)** Plastid gene content of the Apicomplexa and free-living relatives. Presence is denoted by filled circles; variable presence between species within a clade is denoted by half-filled circles; dotted circles represent possible gene remnants (pseudogenes or highly-divergent sequences). **(c)** The corallicola plastid genome, compared to those of *V. brassicaformis* (free-living, photosynthetic) and *T. gondii* (obligate parasite). Gene colors correspond to gene categories in panel (b). Purple bars denote tRNAs. Orange shading link regions of genomic synteny. Genomes are shown at same scale; only select portions of the *V. brassicaformis* plastid genome are shown.

The corallicola plastid genome shares a curious combination of similarities with both the apicoplast, and its photosynthetic relatives. At 46 kb, it was intermediate in size, having lost a substantial number of genes including those encoding photosystems (Fig. 3b). These genes are essential for photosynthesis, and are still plastid-encoded in chromerids (Janouškovec et al. 2010), strongly suggesting that corallicola is non-photosynthetic. However, corallicola also retained a number of other genes that were lost in known apicoplasts. These include 5S rRNA and two genes encoding proteins that interact with it, *rpl5,* and *rps13* (Dontsova & Dinman 2005). Most remarkable was the presence of *chlL*, *chlN*, *chlB*, and *acsF,* four genes involved in chlorophyll biosynthesis (Fig. S4). These are the only four genes in this pathway that the ancestral plastid would have encoded, and phylogenetically, they grouped with homologues from *V. brassicaformis*, indicating that they do derive from the photosynthetic apicomplexan ancestor and were not the result of horizontal acquisition (Fig. S3b,c). These results provide a window into the evolutionary process, showing that apicoplast genome reduction likely occurred in a stepwise manner, whereby all photosystem genes were first lost from the common ancestor with chromerids, followed by the loss of chlorophyll biosynthesis genes in the parasitic apicomplexans (Fig. 3c).

The retention of *chlL*, *chlN*, *chlB*, and *acsF* in the face of otherwise severe gene loss indicates that these genes remain under strong selection. What function underlies this selection is less obvious: corallicola cells were unpigmented and non-autofluorescent (at >550 nm, using 450–500 nm excitation) and, considering the absence of photosystem genes in the plastid genome, are likely non-photosynthetic. Chlorophyll itself has no natural biological function outside of photosynthesis, so corallicola must have evolved a novel use for either chlorophyll or its closely-related precursors or derivatives. These molecules, however, generally function in light harvesting, which would be highly destructive to cellular integrity without the coupling of the resulting high-energy compounds to photosynthesis. Other possibilities are functions in light-sensing, photo-quenching, or heme synthesis regulation, but these too leave open the question of what the cell would do with the high-energy end products. Moreover, we detected corallicola in sun coral (*Tubastrea* sp.) and black coral (order Antipatharia), both of which are typically considered non-photosynthetic corals (Fig. 1c), further suggesting that light harvesting is not of primary importance.

Whatever the function of these genes may be, phylogenetic analyses suggest that corallicola may not be the only apicomplexan lineage to retain them. Whole-plastid genome trees placed corallicola at the base of the Apicomplexa (Fig. S3b; consistent with 16S rRNA analyses [Janouškovec et al. 2012; Mathur et al. 2018]), which was also the most parsimonious with respect to gene content (Fig. 3b). However, nuclear rRNA phylogenies placed corallicola deeply within the parasitic apicomplexans, as sister to Coccidia (Fig. 3a, Toller et al. 2002). The mitochondrial genome phylogeny agreed with this (Fig. S3a), as did plastid gene synteny, amino acid identity, the use of UGA as a tryptophan codon, and single-gene phylogenies of several plastid genes (Fig. S5a,b). There is no straightforward explanation for this incongruence, but the lack of data for deep-branching apicomplexan plastids makes it more likely that the plastid phylogeny is misleading, and that continued sampling will yield other lineages where chlorophyll biosynthesis is retained.

Most reef-dwelling corals are photosynthetic by virtue of symbiosis with *Symbiodinium.* While the nature of corallicola’s interaction with corals almost certainly differs in this regard, there is also no evidence that it is pathogenic. Corallicola and *Symbiodinium* have non-overlapping localizations within host tissue (Fig. 2). Sampled corals were almost exclusively in good health (Table S1) and the presence of corallicola did not appear to be detrimental or correlate with host disease. We speculate that if corallicola does induce pathology, its negative impacts are probably minor, strain-specific, or arise opportunistically. Elucidation of the corallicola life cycle may help assess its impact on coral health and its possible role in reef ecosystems. Upton and Peters (1986) described *G. cylindrus* sporozoites and oocysts in coral tissues, but left unanswered the existence of life stages outside corals, perhaps in another host. Kirk et al. (2013) found that the planulae (larvae) of brooding species already harboured Type-N, indicating vertical transmission, whereas the gametes of broadcast spawning corals did not, suggesting horizontal/environmental acquisition in these species. The potential for mixed modes of transmission contingent on host traits is similarly reported for *Symbiodinium* (Baird et al. 2009; Quigley et al. 2017).

Corals are a diverse group of cnidarians found across temperate and tropical oceans, and are fundamental to the building of coral reef ecosystems. In recent years, there have been alarming losses of healthy reefs worldwide due to stressors such as climate change and pollution. Changes in the microbiome are associated with coral stress and disease (Meyer et al. 2014; Glasl et al. 2016), and rising temperatures can result in coral bleaching due to the expulsion of *Symbiodinium* (Davy et al. 2012). Therefore, understanding the intricate symbiotic relationships between corals and their microbes is crucial in the effort to decipher the processes behind reef degradation. The identity and nature of coral-microbe associations, outside of *Symbiodinium*, remain poorly characterized (Hernandez-Agreda et al. 2017; Ainsworth et al. 2017). Here, we show that diverse anthozoans are ubiquitously colonized by members of a single apicomplexan lineage. Our results indicate that corallicola, like *Symbiodinium*, is a core coral symbiont, but one which represents an unexplored component of the coral microbiome. The discovery of corallicola and its unusual characteristics has implications for both the understanding of coral biology and the evolution of parasitism.

## Methods and Materials

### Microbial community survey of wild corals

Corals were collected from several locations in Curaçao in April 2015, under the collecting permits of the Dutch Antillean Government (Government reference: 2012/ 48584) provided to the CARMABI Foundation (CITES Institution code AN001) (Table S1). Whole samples, including skeleton and tissue, were homogenized using mortar and pestle, and DNA was extracted with the RNA PowerSoil Total RNA Isolation Kit coupled with the DNA Elution Accessory Kit (MoBio). DNA concentration was quantified on a Qubit 2.0 Fluorometer (Thermo Fisher Scientific Inc.).

Prokaryotic microbiome amplicon preparation and sequencing was performed by the Integrated Microbiome Resource facility at the Centre for Comparative Genomics and Evolutionary Bioinformatics at Dalhousie University. PCR amplification from template DNA was performed in duplicate using high-fidelity Phusion polymerase. A single round of PCR was done using "fusion primers" (Illumina adaptors + indices + specific regions) targeting the V6–V8 region of the Bacteria/Archaea 16S rRNA gene (primer set B969F + BA1406R [Comaeu et al. 2011]; ~440–450 bp fragment) with multiplexing. PCR products were verified visually by running a high-throughput Invitrogen 96-well E-gel. The duplicate amplicons from the same samples were pooled in one plate, then cleaned-up and normalized using the high-throughput Invitrogen SequalPrep 96-well Plate Kit. The samples were then pooled to make one library which was then quantified fluorometrically before sequencing on an Illumina MiSeq using a 300 bp paired-end read design.

Eukaryotic microbiome amplicons were prepared using PCR with high-fidelity Phusion polymerase (Thermo Fisher Scientific Inc.), using primers that target the V4 region of 18S rRNA gene, but which exclude metazoan sequences (UnonMetaF 5'-GTGCCAGCAGCCGCG-3', UnonMetaR 5'-TTTAAGTTTCAGCCTTGCG-3') (Bower et al. 2004). PCR was performed using the following protocol: 30s at 98°C, followed by 35 cycles each consisting of 10 s at 98 °C, 30 s at 51.1°C and 1 min at 72°C, ending with 5 min at 72°C. PCR products were visually inspected for successful amplification using gel electrophoresis with 1% agarose gels. PCR products were then cleaned using the QIAquick PCR Purification Kit (Qiagen) and quantified on a Qubit 2.0 Fluorometer. Amplicon sequencing was performed by the Integrated Microbiome Resource facility at the Centre for Comparative Genomics and Evolutionary Bioinformatics at Dalhousie University, as above, but using the Eukaryote-specific primer set E572F + E1009R (~440 bp fragment [Comaeu et al. 2011]).

Amplicon reads were processed (dereplication, chimera detection, and singleton removal) using VSEARCH (Rognes et al. 2016). Operational taxonomic units (OTUs) were clustered at 97% identity using VSEARCH and analyzed using Qiime v1.9.1 (Caporaso et al. 2010). The taxonomic identity of each OTU was assigned based on the SILVA 128.1 database (Quast et al. 2013), modified to include the SSU of the coral skeleton boring algae *Ostreobium queketii* (del Campo et al. 2017), using the assign_taxonomy function in Qiime. OTUs that were “unassigned” were inspected using BLAST against the GenBank nr database and manually reassigned to the closest hit if possible. OTUs represented by fewer than four reads were removed, as were OTUs that were identified as metazoan 18S rDNA or mitochondria. Samples with fewer than 1,500 reads were excluded from downstream analysis. In total, 863,280 reads (avg. 20,554 per sample) were obtained in the eukaryotic 18S rDNA dataset after filtering. For the prokaryotic 16S rDNA dataset, a total of 254,611 reads (8,780 per sample) were obtained after filtering. Due to specificity of the 16S primer set used, *Symbiodinium* plastid 16S rDNA sequences were generally not amplified (Table S2). Differences in detecting ARL-V and Type-N from the same samples may be due to differential read depth between the eukaryotic and prokaryotic datasets.

### Metagenome database mining

Thirty one coral-derived metagenomic and metatranscriptomic assemblies were retrieved from the Joint Genome Institute Integrated Microbial Genomes and Microbiomes (JGI IMG/M) database (Table S3). These were screened for the presence of the ARL-V 16S rRNA gene (DQ200412) and the 18S rRNA genes of Type-N (AF238264), *C. velia* (NC_029806), and *V. brassicaformis* (HM245049) by BLASTn searches. To retrieve Corallicola plastid and mitochondrial sequences, protein-encoding genes of *V. brassicaformis* (HM222968, Flegontov et al. 2015), *C. velia* (HM222967), *T. gondii* (U87145), and *P. falciparum* (LN999985, AY282930) plastids and mitochondria were translated and searched against the datasets with tBLASTn. All hits were manually inspected by BLAST against the GenBank nr database to verify that they correspond to apicomplexan-related organisms. Hits that most closely matched to *Symbiodinium* spp. and *Ostreobium* spp. sequences were discarded. Hits were assembled into longer contigs using Geneious R9 (Biomatters Ltd.). Assembled sequences were deposited in GenBank (see *Data availability*).

### Aquarium coral survey

Coral samples were purchased from Aquariums West (Vancouver, BC, Canada) or online from Canada Corals Inc. (Mississauga, ON, Canada) and Fragbox Corals (Toronto, ON, Canada) (Table S1). Identification was based on morphology and/or vendor labels. Corals were thoroughly rinsed with salt water (Instant Ocean Reef Crystals Salt Mix) and cut into smaller pieces containing at least 1 polyp or, for larger specimens, a portion containing skeleton, tissue, and part of the oral disc. Samples were homogenized using a mortar and pestle, and DNA was extracted with the DNeasy PowerBiofilm Kit (Qiagen) according to manufacturer’s instructions. Samples were screened by PCR for the presence of the Type-N 18S rRNA gene using primers 18N-F2 (5’-TAGGAATCTAAACCTCTTCCA-3’) and 18N-R1 (5’-CAGGAACAAGGGTTCCCGACC-3’) (Toller et al. 2002). PCR was performed with Phusion DNA polymerase (Thermo Fisher Scientific Inc.) using 32 cycles of amplification (98°C for 8 s, 60.5°C for 15 s and 72°C for 30 s) after an initial incubation for 30 s at 98°C. Selected amplicons were Sanger sequenced to verify that the Type-N 18S rRNA gene was correctly amplified.

### Fluorescence microscopy

Corallicola was visualized in the green mushroom coral (Corallimorpharia, Discosomatidae, *Rhodactis* sp.), which lacks a calcium carbonate skeleton, has a large polyp structure, and is amenable to tissue fixation with the following method. Dissected tissues were placed directly in Carnoy’s solution (6:3:1 ethanol:chloroform:acetic acid) and soaked overnight. Tissues were then washed with 80% (v/v) ethanol for 3 × 10 min, and in bleaching solution (80% v/v ethanol, 6% v/v H_2_O_2_) for 2 × 10 min. Samples were left in bleaching solution for 7 days, with replacement of solution every other day. After bleaching, tissues were washed with 100% ethanol for 2 × 10 min, PBSTx (0.3% v/v Triton X-100 in phosphate buffered saline, pH 7.4) for 3 × 10 min, and with hybridization buffer (0.9 M NaCl, 20 mM Tris-HCl, 30% v/v formamide, 0.01% w/v sodium dodecyl sulfate) for 3 × 10 min. Fluorescent *in-situ* hybridization was carried out by incubating tissues in hybridization buffer with DAPI DNA stain (0.01 mg/ml) and fluorescent labeled DNA probes (0.1 µM), overnight in the dark with agitation. Samples were then washed with PBSTx for 3 × 10 min, placed on glass slides with ProLong Diamond Antifade Mountant (Thermo Fisher Scientific Inc.), and visualized on a Zeiss Axioplan 2 microscope. The following probes were used: wk16P (5’-CTGCGCATATAAGGAATTAC-3’) with 5’ Texas Red-X label, targeting Type-N 18S rRNA; wk17P (5’-TCAGAAGAAAGTCAAAAACG-3’) with 5’ Alexa Fluor 532 label, targeting ARL-V 16S rRNA; wk18P (5’-GCCTTCCCACATCGTTT-3’) with 5’ Texas Red-X label, targeting Gammaproteobacteria as a control.

### Electron microscopy

Mesenterial filaments from *Rhodactis* sp. were fixed in 2.5% glutaraldehyde in 35 ppt salt water (Instant Ocean Reef Crystals Salt Mix), then rinsed 3× in 0.1 M sodium cacodylate buffer and postfixed with 1% osmium tetroxide in 0.1 M sodium cacodylate buffer. Samples were then rinsed 3× in distilled water, dehydrated with successive washes in 30%, 50%, 70%, 90%, 95%, and 3× 100% ethanol, and embeded in Spurr's epoxy resin following infiltration. Resulting blocks were cut into 70 nm sections, and stained using 2% aqueous uranyl acetate (12 min) and 2% aqueous lead citrate (6 min). Sections were viewed under a Hitachi H7600 transmission electron microscope (UBC Bioimaging Facility, Vancouver, Canada).

### Metagenomic sequencing and plastid closing

To enrich for corallicola, the cnidoglandular lobes of *Rhodactis* sp. were removed from mesenterial filaments after soaking in 100% ethanol. DNA was extracted using the DNeasy PowerBiofilm Kit (Qiagen). Two Nextera XT libraries were generated using 1 ng and 10 ng of template DNA, according manufacturer’s instructions (Illumina Inc.). Libraries were sequenced on a single lane of Illumina HiSeq 2500 (The Centre for Applied Genomics, The Hospital for Sick Children, Toronto, Canada), generating 260.8 million paired-end 125 bp reads. Reads were assembled with Megahit 1.1.2 (Li et al. 2016). The mitochondrial genome of the coral and the 18S, 5.8S, and 28S rRNA genes of corallicola were retrieved from the assembly. Reads were mapped using Bowtie2 (Langmead & Salzberg 2012) against mitochondria and plastid contigs assembled from the 31 previous metagenomic datasets (see above). Read coverage was < 1×; therefore, to close the corallicola organelle genomes, gap-spanning regions were PCR amplified with sequence-specific primers and Phusion DNA polymerase (Thermo Fisher Scientific Inc). Amplicons were sequenced by the Sanger method, and resulting reads were assembled in Geneious R9 (Biomatters Ltd.). A predicted 68 bp hairpin region in the plastid genome was unable to be bridged by PCR. The sequence for this region was filled in using reads from the *Cyphastrea* sp. 2 (GOLD Analysis Project ID Ga0126343) metagenome that spanned the gap.

### Phylogenetic analyses

#### Nuclear rRNA gene phylogeny (Fig. 3a)

18S and 28S rRNA sequences were aligned with SINA 1.2.11 (Pruesse et al. 2012), and 5.8S rRNA sequences were aligned with MUSCLE and edited manually in Geneious R9 (Biomatters Ltd.). Alignments of the three genes were concatenated. A maximum likelihood phylogeny was built with the GTR + GAMMA model (1,000 bootstrap replicates) in RAxML 8.2.10 (Stamatakis 2014). A phylogeny based on Bayesian inference was constructed using the GTR + GAMMA model (2 × 10^5^ generation run-time with tree sampling every 200 generations, 0.25 fraction burn-in) in MrBayes 3.2 (Ronquist et al. 2012).

#### Mitochondrial protein phylogeny (Fig. S3a)

The three mito-chondria-encoded genes, *cox1*, *cox3*, and *cob*, were translated and aligned with MUSCLE in Geneious R9. The genes from *V. brassicaformis* were excluded due to their extreme divergence and the resulting long branch in the final tree. Phylogenies were constructed in RAxML 8.2.10 (MtZoa + GAMMA model, 1,000 bootstraps) and MrBayes 3.2 (mixed + GAMMA model, 10^5^ generation run-time with tree sampling every 200 generations, 0.25 fraction burn-in).

#### Plastid protein phylogeny (Fig. S3b)

Nineteen plastid-encoded genes common to apicomplexans, chromerids, and stramenophiles (*rpl2*, *rpl4*, *rpl6*, *rpl14*, *rpl16*, *rps2*, *rps3*, *rps4*, *rps7*, *rps8*, *rps11*, *rps12*, *rps17*, *rps19*, *clpC*, *rpoB*, *rpoC1*, *rpoC2*, and *tufA*) were translated and aligned with MUSCLE in Geneious R9. Split genes were concatenated, and the most conserved copy of duplicated (paralogous) genes were used for this analysis. Genes were translated with Genetic Code 4, as the TGA stop codon codes for tryptophan in chromerid and apicomplexan plastids, including in that of Corallicola. Protein sequences were concatenated and phylogenies were built in RAxML 8.2.10 (cpREV + GAMMA + F model, 1,000 bootstraps), in MrBayes 3.2 (cpREV + GAMMA model, 3 × 10^5^ generation run-time with tree sampling every 500 generations, 0.25 fraction burn-in), and with the neighbor-joining algorithm (Jukes-Cantor model, 1,000 bootstraps). Phylogenies for concatenated ChlN, ChlB, and ChlL proteins, and for AcsF (Fig. S3c), were also generated as above with RAxML 8.2.10 and with MrBayes 3.2 (cpREV + GAMMA model, 10^5^ generation run-time with tree sampling every 5200 generations, 0.25 fraction burn-in).

#### Plastid, single gene analysis (Fig. S5b)

Phylogenetic trees were constructed for each of the above 19 proteins, as well as for SufB, Rps5, Rpl11, Rpl36, and the 16S and 23S rRNA genes. MUSCLE and SINA were used to align proteins and rRNAs, respectively. Trees were built using RAxML 8.2.10 (cpREV + GAMMA + F and GTR + GAMMA models, 500 bootstraps), MrBayes 3.2 (cpREV + GAMMA model and GTR + GAMMA and Poisson + GAMMA models, tree sampling every 200 generations for 10^5^ generations, 0.25 fraction burn-in), neighbor-joining (Jukes-Cantor model, 1,000 bootstraps), and maximum parsimony (all sites, 100 bootstraps) in MEGA 7 (Kumar et al. 2016).

Accession numbers of sequences used in this study are listed in Table S5.

#### Other methods

Coral phylogeny (Fig. 1c) was based on the established topology of the clades (Kayal et al. 2013, Zapata et al. 2015) and on published analyses that use molecular clocks and fossils to date the age of clade divergences (Simpson et al. 2011, Stolarski et al. 2011, Park et al. 2012, Baliński et al. 2012).

Pairwise dN values (number of nonsynonymous substitutions per nonsynonymous site) were calculated from translation-aligned nucleotide sequences using codeml from the PAML package (Yang 2007). The following settings were used: Seqtype = 1: codons, alpha = 0 (fixed), Small_Diff = 5e-07, model = 0: one w, runmode = -2, clock = 0, Mgene = 0, CodonFreq = 2: F3X4, estFreq = 0, fix_blength = 0, optimization method = 0, icode = 3: mold mt.

### Data availability

The following are deposited in GenBank: The *Rhodactis* sp. wkC1 mitochondrial genome (accession no. MH320096); Corallicola 18S/5.8S/28S rRNA genes from *Rhodactis* sp. wkC1 (MH304758, MH304759), *Orbicella* sp. TRC (MH304760), *Cyphastrea* sp. 2 (MH304761); Corallicola mitochondrial genomes from *Rhodactis* sp. wkC1 (MH320093), *Orbicella* sp. 8CC (MH320094), *Cyphastrea* sp. 2 (MH320095); Corallicola plastid genome from *Rhodactis* sp. wkC1 (MH304845).

The 18S and 16S rRNA gene amplicon reads are deposited in the NCBI Sequence Read Archive (PRJNA482746).

## Acknowledgements

We thank Charlotte Zwimpfer and Bradford Ross for assistance in sample processing and electron microscopy. This work was funded by a grant from the Canadian Institutes for Health Research (MOP-42517). W.K.K. was supported by a Natural Sciences and Engineering Research Council of Canada Fellowship (PDF-502457-2017) and a Killam Post-doctoral Research Fellowship, J.dC was supported by a grant from the Tula Foundation to the Centre for Microbial Biodiversity and Evolution and the Marie Curie International Outgoing Fellowship FP7-PEOPLE-2012-IOF - 331450 CAARL.

## Author Contributions

W.K.K., J.dC., and P.J.K. designed the study. W.K.K., J.dC., M.J.A.V., and P.J.K. obtained samples. J.dC. and V.M. conducted microbial community analyses. W.K.K. performed all other analyses. W.K.K. and P.J.K. wrote the manuscript with input from all authors.

## Supplementary Figures & Tables

**Figure S1.** Mitochondrial genomes of corallicola. Names denote the host coral from which the genomes were retrieved. The three mitochondria-encoded genes are shown in blue. Clockwise tick marks denote 1,000 bp. It is unclear whether the genomes are circular (as depicted), or tandem linear.

**Figure S2.** Transmission electron micrographs of darkly-staining organelles in corallicola, showing distinctive internal structures. Structure and orientations (sagittal and transverse sections illustrated at top) were inferred from viewing multiple organelles from several cells.

**Figure S3. (a)** Phylogenetic placement of corallicola, based on mitochondria-encoded proteins. **(b)** Phylogenetic placement of corallicola, based on plastid-encoded proteins. **(c)** Phylogenetic placement of corallicola’s chlorophyll biosynthesis proteins (concatenation of proteins ChlL, ChlN, and ChlB at left; AcsF at right). All phylogenetic trees shown were produced with the maximum-likelihood algorithm; values at nodes denote maximum-likelihood bootstrap support percentages (n = 1,000 replicates) and Bayesian posterior probabilities (see Methods and Materials).

**Figure S4.** Tetrapyrrole and chlorophyll biosynthesis pathways, showing the function of genes retained in the corallicola plastid genome (*acsF*, *chlL*, *chlN*, and *chlB*; highlighted). All enzymatic steps depicted here are inferred to occur within the apicomplexan plastid (Koreny et al. 2011).

**Figure S5. (a)** Pairwise amino acid identities and dN values support a close relationship between corallicola and the Coccidia. **(b)** Phylogenetic analysis of single plastid genes/proteins to test alternative topologies of corallicola placement. Results vary by gene and methodology: while most plastid genes show a basal placement for corallicola, a few support the grouping of corallicola within the Apicomplexa. Tree construction methods are indicated at top, with the model of evolution in parentheses: NJ, neighbour-joining; MP, maximum parsimony; ML, maximum-likelihood. A dash indicates a lack of support for either topology.

**Table S1.** Coral samples used in this study.

**Table S2.** Eukaryote & prokaryote OTU tables and representative sequences.

**Table S3.** Summary of metagenomic/metatranscriptomic dataset survey results, with list of contigs corresponding to the corallicola plastid genome.

**Table S4.** Corallicola prevalence from previous studies.

**Table S5.** Nucleotide sequences used in this study.

